# Early Differentiation Signatures in Human Induced Pluripotent Stem Cells Determined by Non-Targeted Metabolomics Analysis

**DOI:** 10.1101/2023.03.02.530741

**Authors:** Rodi Abdalkader, Romanas Chaleckis, Takuya Fujita

## Abstract

Human induced pluripotent stem cells (hiPSCs) possess immense potential as a valuable source for the generation of a wide variety of human cells yet monitoring the early cell differentiation towards a specific lineage remains challenging. In this study, we employed a non-targeted metabolomic analysis technique to analyze the extracellular metabolites present in samples as small as one microliter. The hiPSCs were subjected to differentiation by initiating culture under the basal medium E6 in combination with chemical inhibitors that have been previously reported to direct differentiation towards the ectodermal lineage such as Wnt/β- catenin and TGF-β kinase/activin receptor alone or in combination with bFGF, and the inhibition of glycogen kinase 3 (GSK-3), which is commonly used for the diversion of hiPSCs towards mesodermal lineage. At 0 hr and 48 hrs 107 metabolites were identified, including biologically relevant metabolites such as lactic acid, pyruvic acid, and amino acids. By determining the expression of the pluripotency marker OCT3/4, we were able to correlate the differentiation status of cells with the shifted metabolites. The group of cells undergoing ectodermal differentiation showed a greater reduction in OCT3/4 expression. Moreover, metabolites such as pyruvic acid and kynurenine showed dramatic change under ectodermal differentiation conditions where pyruvic acid consumption increased 1-2-folds, while kynurenine secretion decreased 2-folds. Further metabolite analysis uncovered a group of metabolites specifically associated with ectodermal lineage, highlighting the potential of our findings to determine the characteristics of hiPSCs during cell differentiation, particularly under ectodermal lineage conditions.

**Graphical abstract:** 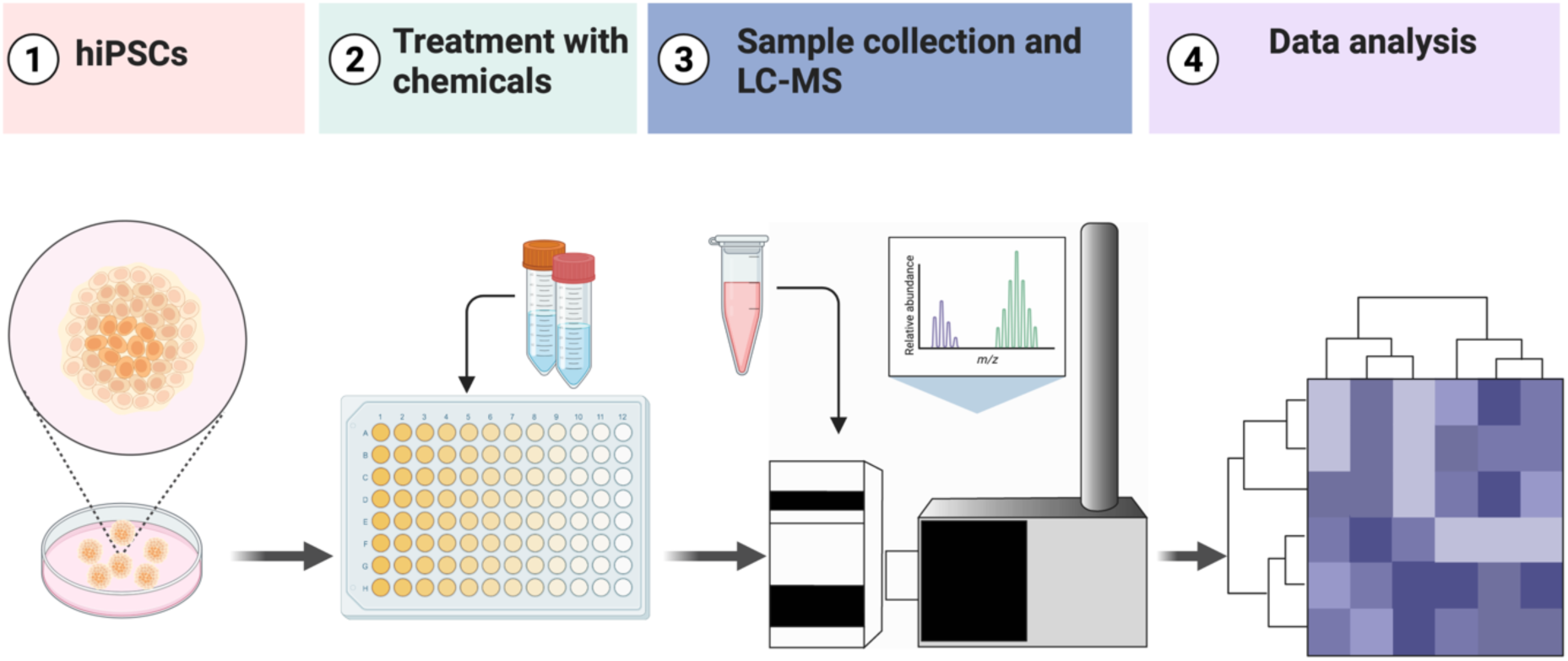

## 1. Introduction

Human pluripotent stem cells (hPSCs) are divided into human embryonic stem cells (hESCs), which are generated from the inner cellular mass of the blastocyst of pre-implantation embryos^1^, and into human induced pluripotent stem cells (hiPSCs), which are directly generated from somatic cells upon the introduction of four transcription factors (Oct3/4, Sox2, Klf4, and Myc)^2^. hPSCs can self-renew or divert into the three germ layers: ectoderm, mesoderm, and endoderm under specific conditions to form functional cells such as liver cells ^3, 4^, heart cells ^5^, eye cells ^6, 7^ and others ^8^. Thus, hPSCs provide a valuable source for the generation of human functional cells to be used in various applications such as in cell-based therapies as well as in the construction of *in vitro* models.

The initial differentiation of hPSCs into specific cell lineages can be determined by evaluating the gene expression and protein expression of the three-lineage markers (ectoderm, mesoderm, and endoderm) through quantitative real-time polymerase chain reaction (PCR) for gene expression, western blotting, immunofluorescence staining, or ELISA for protein detection^9^. Although these methods provide useful information, they are invasive as they require the destruction of cells to access the intercellular nucleic acids and proteins. Furthermore, these methods are time-consuming as they entail multiple steps such as the addition of different antibodies and labeling agents and can also be costly. For the purpose of producing hPSCs-derived functional cells on a large scale, non-invasive, simple, quick, and comprehensive analytical methods are desirable for predicting and monitoring hPSCs lineage in the early stage of differentiation to minimize waste of time and materials during cell production.

Metabolome can provide relevant information on cell characteristics^10^. One of the metabolomics techniques, LC-MS-based non-targeted metabolomics, is capable of detecting hundreds of metabolites in small volumes through a relatively straightforward sample preparation process^11, 12, 13^. In previous studies, we successfully used this technique to determine the metabolites in the cell culture medium (CCM) of liver cells (HepG2) and corneal epithelial cells (HCE-T) grown in microfluidic devices by collecting only a small volume at different time points ^14, 15^. Currently, there is no established method for evaluating the lineage deviation of hPSCs during differentiation by profiling extracellular metabolites. Thus, the implementation of LC-MS-based non-targeted metabolomics approach presents a non-invasive solution to predict the differentiation status of hPSCs.

In this study, the early differentiation of hiPSCs was investigated by using chemical inhibitors that have previously been reported to induce differentiation towards the ectodermal and mesodermal lineages. CCM was collected at 0 hr and 48 hrs, and the levels of extracellular metabolites were obtained through LC-MS-based non-targeted metabolomics measurements. The resulting shifts in metabolites were then evaluated in relation to the pluripotency status of the hiPSCs.

## 2. Materials and methods

### 2.1. Culturing human induced pluripotent stem cells

The human induced pluripotent stem cells 585A1 were purchased from Riken cell bank (RIKEN BRC #HPS0354)^16^ and was used following the approval of the Ethics Committee of Ritsumeikan University. Prior to inducing differentiation, the cell culture dish was coated with matrigel and incubated at 37 °C for 30 min. Cultured cells were washed with DPBS and treated with TryPLE Express (Thermo Fisher Scientific, Inc., Waltham, MA, USA) at 37 °C for 5 min, followed by the addition of mTeSR^TM^ plus medium (mTeSR plus) (STEMCELL Technologies) and transfer of the cell suspension into a 15-mL tube. Cells were centrifuged at 200 × *g* for 3 min, and the supernatant was removed and resuspended in mTeSR plus medium supplemented with 10 µM Y27632 (Wako, Osaka, Japan), and plated in 96-well plate at a density of 1×10^3^ cells per well on a matrigel (Thermo Fisher Scientific, Inc., Waltham, MA, USA) coated culture dish, and cultured for 24 h in a humidified incubator at 37 °C with 5% CO_2_. The culture medium was replaced with mTeSR plus basal medium daily while skipping weekends according to the manufacturer’s instruction.

### 2.2. Differentiation induction, sample collection, and cell counting

At 30-40% confluency, the medium was changed to E6 basal medium (Thermo Fisher Scientific, Inc., Waltham, MA, USA) with chemical inhibitors and bFGF growth factor (as listed in Table 1). A sample of the culture medium (1 microliter) was taken at 0 hr and 48 hrs and stored at-80°C. The sample was then dried in a vacuum rotor (PV-1200; Wako, Osaka, Japan) for 20 minutes. To determine cell count, the remaining culture medium was removed, and the cell count was measured using a cell count normalization kit according to the manufacturer’s instructions (Dojindo Laboratories, Japan).

**Table. 1.** The cell culturing conditions of hiPSCs

| Medium | Additives | Working conc. |
| --- | --- | --- |
| mTeSR plus |  |  |
| E6 | DMSO | 0.2 % v/v |
| E6 | bFGF | 50 ng mL<sup>-1</sup> |
| E6 | IWR-1 endo | 2.5 μM |
| E6 | A83-01 | 2.5 μM |
| E6 | IWR-1 endo_A83-01_bFGF | 2.5 μM_2.5 μM_50 ng mL<sup>-1</sup> |
| E6 | IWP-2 | 10 μM |
| E6 | SB-505124 | 10 μM |
| E6 | IWP-2_SB-505124_bFGF | 10 μM_10 μM_50 ng mL<sup>-1</sup> |
| E6 | Y-27632 | 10 μM |
| E6 | CHIR99021_Y-27632_bFGF | 10 μM_10 μM_50 ng mL<sup>-1</sup> |
| E6 | IWP-2_A83-01_bFGF | 10 μM_2.5 μM_50 ng mL<sup>-1</sup> |
| E6 | IWR-1 endo_SB-505124_bFGF | 2.5 μM_10 μM_50 ng mL<sup>-1</sup> |

### 2.3. Non-targeted metabolomic using LSMS analysis

Tubes containing one microliter of dried samples were thawed and 150 μL of water and acetonitrile (3:7, v/v) was added, containing three technical internal standards (tISs): 0.1µM CHES, 0.1µM HEPES, and 0.2µM PIPES. After resuspension, all samples were centrifuged at room temperature for 1 minute at 1000 *g*. A quality control (QC) sample was prepared by pooling an aliquot from each sample. Next, 40 μL of the supernatant was transferred to a 96- well 0.2 mL PCR plate (PCR-96-MJ; BMBio, Tokyo, Japan). The plate was sealed with a pierceable seal (4titude; Wotton, UK) for 3 seconds at 180°C by using a plate sealer (BioRad PX-1; CA, USA) and maintained at 10°C during the LC-MS measurements. The LC-MS method has been described previously ^17, 18, 19^. The injection volumes were 7 μL in negative and 3 μL in positive ionization mode. In brief, metabolite separation was achieved on an Agilent 1290 Infinity II system by using SeQuant ZIC-pHILIC (Merck, Darmstadt, Germany) column by using a gradient of acetonitrile and 5 mM ammonium acetate in water (pH=9.3). Data were acquired on an Agilent 6550 Q-TOF-MS system with a mass range of 40−1200 m/z in all ion fragmentation mode, including three sequential experiments at alternating collision energies: full scan at 0 eV, followed by MS/MS scans at 10 eV and 30 eV, with a data acquisition rate of 4 scans/s. Data were converted to mzML format using Proteowizard and processed using MS-DIAL version 4.80 ^20, 21, 22^. An in-house library containing accurate masses (AM) and retention times (RT) for 609 compounds obtained from chemical standards was used to annotate the detected compounds. Peak areas exported from MS-DIAL were used for metabolites semi-quantification. Only metabolites with a coefficient of variation (CV) less than 30% in the QC samples were used for further analysis.

### 2.4. Immunofluorescence and microscopy imaging

For immunostaining, cells were fixed with 4% paraformaldehyde in PBS for 25 min at 25°C and then permeabilized with 0.5% Triton X-100 in PBS overnight at 4°C. Subsequently, the cells were blocked with blocking buffer [5% bovine serum albumin, 0.1% (v/v) Tween-20] at room temperature for 90 min and then incubated at 4°C overnight with the primary antibodies in blocking buffer (Alexa Fluor 594 anti-OCT3/4, 1:200 v/v; Santa Cruz, sc-5279). The cells were then washed and incubated at 25°C for 60 min with the secondary antibody (Alexa Fluor 594 Goat anti-mouse IgG, 1:500 v/v; Santa Cruz). Finally, the cells were washed and incubated with anti-phalloidin for F-actin using ActinGreenTM 488 (Invitrogen-2277811) at 25°C for 60 min. For imaging, we used a fluorescence microscope (KEYENCE, Tokyo, Japan) for the acquirement of cell images. Cell Profiler software (Version 3.1.8; Broad Institute of Harvard and MIT, USA) was used to determine OCT3/4 expression, using a pipeline that identifies primary nucleus objects based on their diameter and fluorescence intensity of the nucleus stain, using either auto or Otsu segmentation ^23^.

### 2.5. Statistical analysis and data visualization

Independent biological triplicate samples were used for the study. The statistical analysis was performed using the Dunnett’s test and Tukey’s comparison test through GraphPad Prism 8 (GraphPad Software, La Jolla, California, USA). Principal component analysis visualization was performed using MetaboAnalyst platform ^24^. Venn diagrams were generated using Orange 3 software (Version 3.23.1) developed by the Bioinformatics Laboratory at the Faculty of Computer and Information Science, University of Ljubljana, Slovenia. Data visualization was done using GraphPad Prism 8 and R Studio with the ggplot library. The graphical abstract was created using BioRender.com.

## 3. Results

### 3.1. The determination of cells counts and the expression of the pluripotent marker (OCT3/4)

We conducted Hoechst staining at the end of the extracellular metabolites collection process, which lasted for 48 hrs. The purpose of this staining was to accurately determine the cell count. The results of the analysis showed no notable change in the mean fluorescence intensity of Hoechst across all the samples, as depicted in Figure 1A.

**Figure 1.**
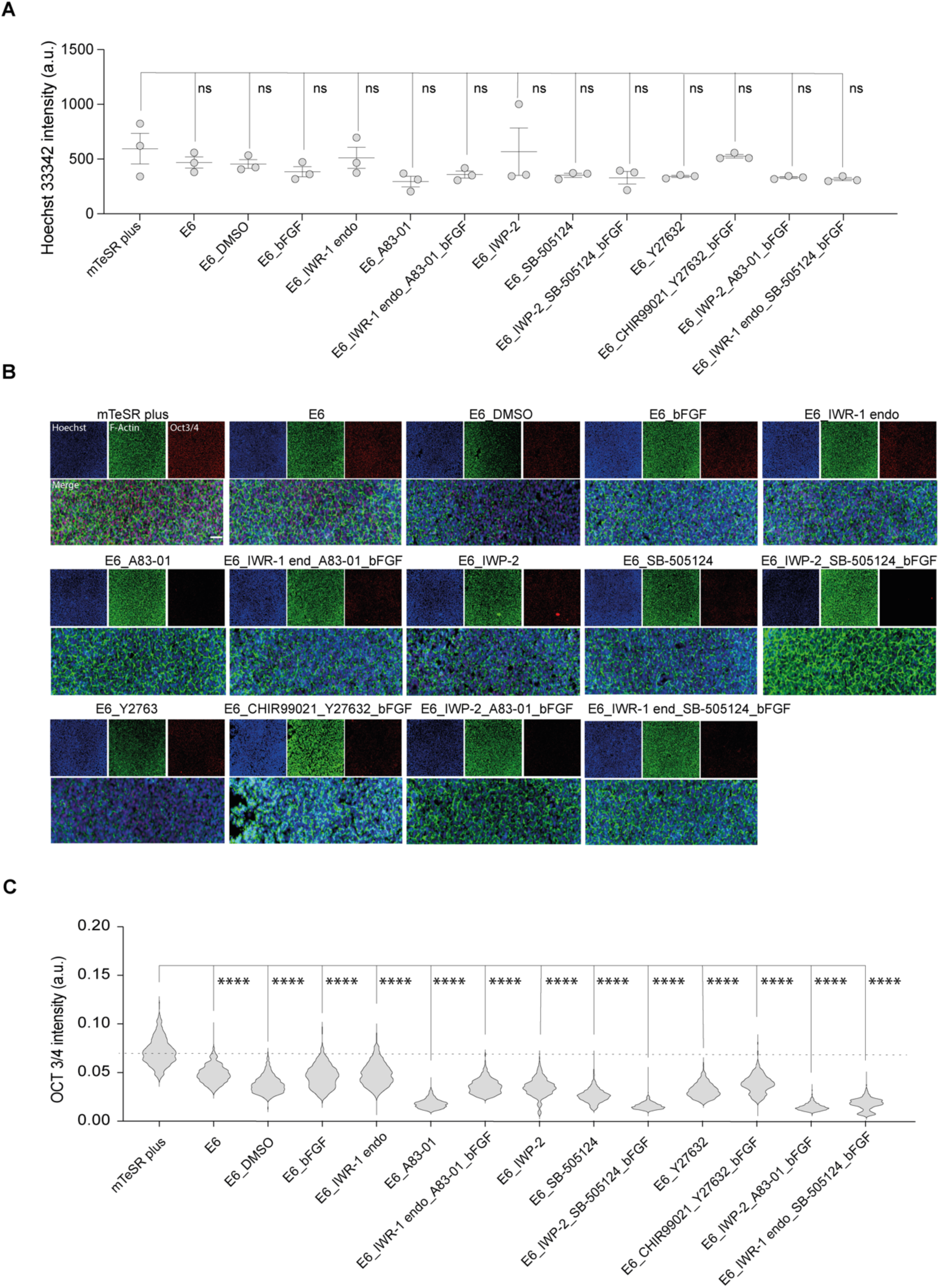
The determination of cells characteristics after samples collection. (A) Cell nucleus count determination after 48 hrs of treatments. Cells were treated with the differentiation induction medium containing the chemical inhibitors with or without bFGF for 48 hrs. And then cells were stained with the Hoechst 33342 for the determination of cells count. Data are presented in biological triplicates as means ± S.E.M. The *p*-values were determined using Dunnett’s test; (ns: not significant). (B) Fluorescent micrograph images indicating the expression of pluripotency marker (OCT3/4) in red, nucleus in blue, and actin fibers (F-actin) in green. Scale bar, 50 μm. (C) Single cells immunofluorescence analysis of OCT3/4. Data are represented in the violin plot in which the median of each group is indicated with a scatter line (25^th^ to 75^th^ interquartile range). The *p*-values were determined using Dunnett’s test. ****: *p* <0.0001.

In order to assess the differentiation status of cells subjected to chemical induction, we performed immunofluorescent staining for the pluripotency protein marker OCT3/4, which is a well-established indicator of pluripotency in cells. The results showed a marked reduction in OCT3/4 expression in all samples treated with E6 differentiation basal medium compared to those maintained in mTeSR plus hiPSCs medium (Figure 1B). Further analysis revealed that the greatest reduction in OCT3/4 expression was seen in groups treated with A83-01 alone, as well as in groups treated with a combination of IWP-2_ SB505124_ bFGF, IWP-2_ A83-01_ bFGF, and IWR-1 endo_ SB505124_ bFGF (Figure 1C).

### 3.2. Measurement of extracellular metabolites and their shifts after 48 hrs

We could annotate 117 metabolites at Metabolomics Standard Initiative annotation level 1 ^25^. Peak areas were used for metabolite semi-quantification. The CVs of the two tIS in the QC samples were <18% and <14% in negative and positive ionization modes; and in the study samples <17% and <12% in negative and positive ionization modes, respectively (HEPES was present at high concentration in cell culture medium resulting in a saturated signal and therefore was not used as tIS). Annotated metabolites AM and RT differences were <15 ppm and <0.9 min, respectively. Several metabolites were detected in single or few samples resulting only in low levels in the QC samples and a low (<3) QC average to Blank ratio. In such case, only metabolites with D-Ratio < 30 was kept.

The results of the principal component analysis (PCA) show a change in metabolite levels over time, as indicated by a systematic shift in the sample clusters at 48 hrs compared to 0 hr. Additionally, hiPSC samples in mTeSR plus were distinguishable from the samples that underwent differentiation in E6 medium at both 0 hr and 48 hrs (Figure 2A). The partial least square discrimination analysis (PLS-DA) combined with the variable importance in projection (VIP) was utilized to uncover unique metabolites in the study samples. This analysis showed a significant increase in lactic acid levels, from virtually undetectable at 0 hr to 100-fold higher at 48 hrs, across all samples (Figure 2B). Conversely, the differentiation process led to a decrease of 1-2-fold in pyruvic acid levels. The analysis also indicated correlations between glucose, arginine, and hypoxanthine with pyruvic acid (Figure 2C).

**Figure 2.**
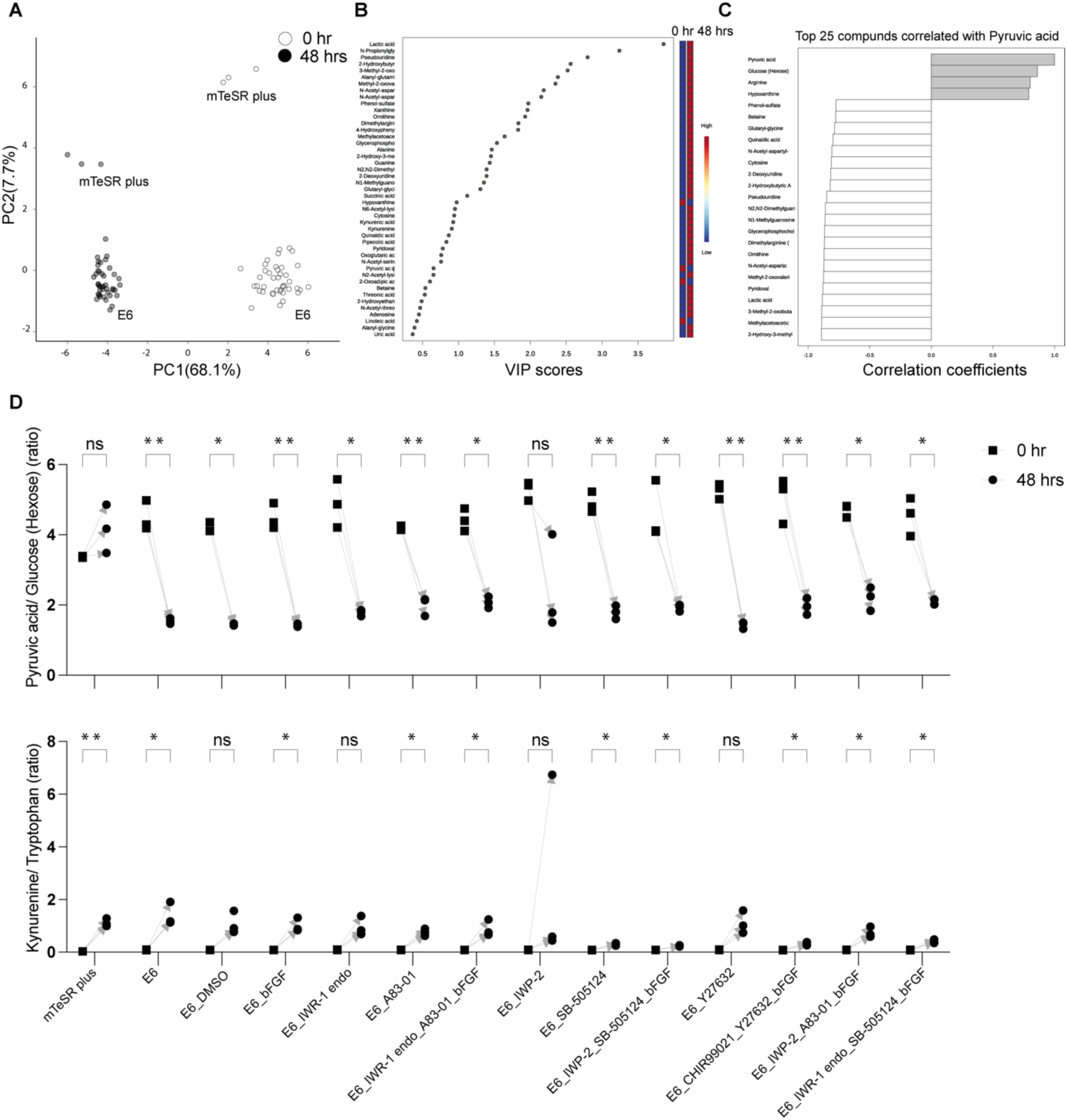
Temporal analysis of extracellular metabolites from the hiPSC-585A1 at 0 hr and 48 hrs. (A) Principal component analysis (PCA) of metabolomics dataset peak areas where samples at 0 hr and 48 hrs are highlighted in different colors. Data are derived from three biological replicates. (B) Variable importance in projection (VIP) derived from the Partial least square discrimination analysis (PLS-DA) for the selection of important metabolites. (C) Pearson correlation analysis of top 25 metabolites with pyruvic acid. (D) Representative examples of selected metabolites were analyzed by calculating their peak area ratios, such as pyruvic acid to glucose and kynurenine to tryptophan. The *p*-values were determined using a paired t-test, with a significance level of * *p* <0.05 and ** *p* <0.01. No significant difference was noted for metabolites with a *p*-value of “ns”.

The hiPSCs cultured under mTeSR plus showed elevated secretion levels of lactic acid. However, no significant changes in pyruvic acid were seen, as determined by the analysis of the raw peak area values and the ratios between pyruvic acid and its glucose precursor. Conversely, the hiPSCs undergoing differentiation in the presence of E6 and chemicals displayed depleted levels of exogenous pyruvic acid and a decrease in kynurenine secretion. The latter was evaluated through the analysis of raw peak area values and the ratios of peak area values between kynurenine and its precursor tryptophan, across several differentiation conditions, including CHIR99021_Y-27632_bFGF, IWP-2, IWP-2_SB-505124_bFGF, IWR- 1 endo_ SB-505124_ bFGF, and SB-505124. (Figure S1) (Figure 2D).

Next, we analyzed the changes in metabolites over 48 hrs by calculating the fold-change (FC) against the initial metabolite raw values at 0 hr (**Supplementary data** table 1). The metabolites were then divided into three categories based on their FC value: Category i (log2 fold change ≤-1), in which metabolites were depleted; Category ii (log2 >-1 and log2 < 1), in which there were no notable secretion or depletion; and Category iii (log2 fold change ≥1), in which nutrient metabolites were secreted. Samples under mTeSR plus medium conditions had 9 metabolites in Category i, 68 metabolites in Category ii, and 39 metabolites in Category iii. The cells grown in basal E6 medium showed 4 metabolites in Category i, 68 metabolites in Category ii, and 45 metabolites in Category iii. To focus on the most related metabolites to cell differentiation, we examined the groups that had the greatest decrease in the pluripotency marker (OCT3/4). These included A83-01 (Category i: 4; Category ii: 72; Category iii: 43), IWP-2_SB505124_bFGF (Category i: 2; Category ii: 63; Category iii: 44), IWP-2_A83- 01_bFGF (Category i: 2; Category ii: 72; Category iii: 43), and IWR-1 endo_SB505124_bFGF (Category i: 10; Category ii: 63; Category iii: 44) (Figure 3) (Figure S2).

**Figure 3.**
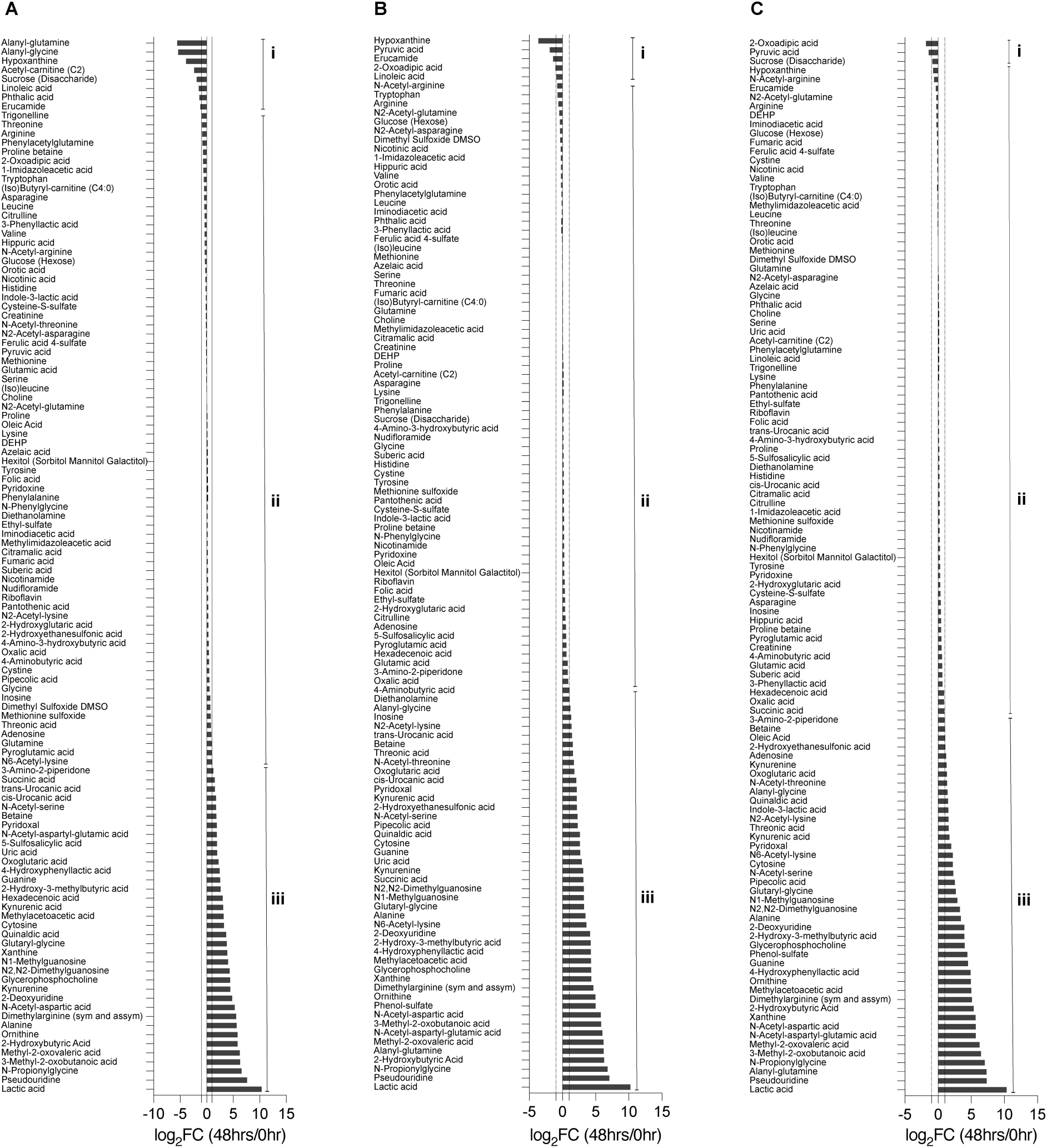
The determination of metabolites changes after 48 hrs. The ratio folds change (log_2_ 48 hrs/0 hr) of metabolites abundances. (A) Under hiPSCs medium (mTeSR plus) conditions. (B) Under the differentiation basal E6 medium conditions. (C) Under the differentiation basal E6 medium conditions in combination with IWP-2 _ SB505124_ bFGF. Category I (log2 fold change ≤-1), where metabolites were depleted, Category ii (log2>-1 and log2 < 1), where metabolites had no notable secretion or depletion, and Category iii (log2 fold change ≥1), in which metabolites were secreted.

To evaluate the impact of various treatments on metabolite changes, metabolites were categorized and presented through Venn diagrams comparing the samples under mTeSR plus and E6 medium, as well as between the chemical treatment groups under the E6 differentiation medium (IWP-2_SB505124_bFGF and IWR-1 endo_SB505124_bFGF) and E6 alone. The comparison between mTeSR plus and E6 medium revealed the presence of 2 common metabolites in Category ii, 7 metabolites found to be higher in mTeSR plus (such as glutamine and pyroglutamic acid), and 2 metabolites specific to E6 medium (such as pyruvic acid and 2- oxoadipic acid) in Category i. In Category iii, there were 34 common metabolites, 5 metabolites that were higher in mTeSR plus (such as glutamine and pyroglutamic acid), and 11 metabolites that were specific to E6 medium (such as inosine) (Figure S3).

In the comparison between the chemical treatment groups under the E6 differentiation medium and E6 alone, there were 2 common metabolites in Category i, with 7 metabolites found to be higher in IWR-1 endo_SB505124_bFGF (such as oleic acid and linoleic acid). In Category iii, there were 38 common metabolites, with 3 metabolites found to be higher in IWP- 2_SB505124_ bFGF (such as indole-3-lactic acid), 4 metabolites found to be higher in IWR-1 endo_SB505124_bFGF (such as oxalic acid, and 5-sulfosalicylic acid), and 5 metabolites found to be higher in E6 alone (such as uric acid, cis/trans-urocanic acid, and inosine) (Figure 4).

**Figure 4.**
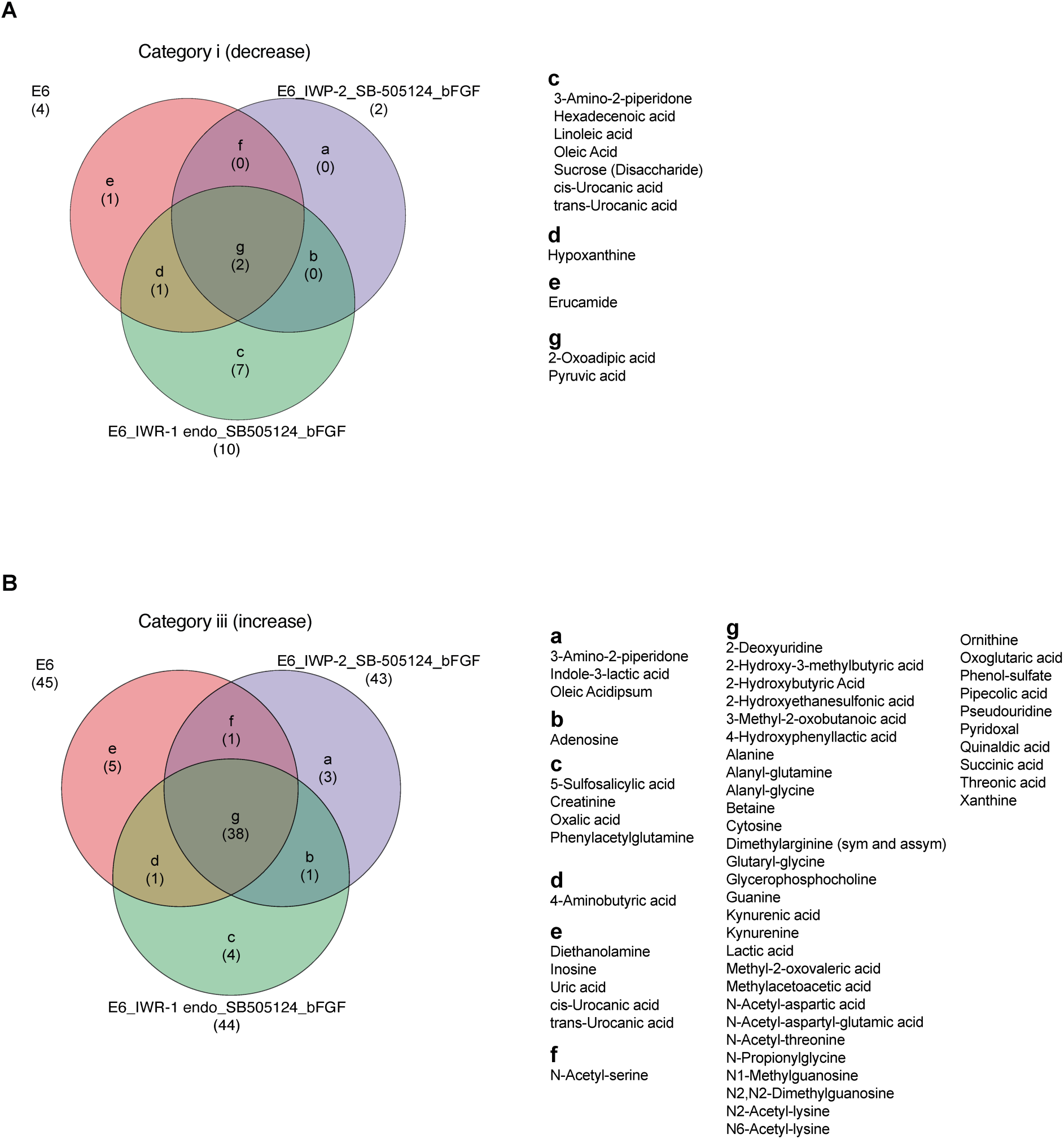
Metabolites signatures under the ectodermal differentiation conditions. Venn diagrams that highlight the differential shifts of metabolites between the differentiation medium E6, E6 in combination with IWP-2 _ SB505124_ bFGF, and E6 in combination with IWR-1 endo _ SB505124_ bFGF. (A) In Category i (log2 fold change ≤-1). (B) In Category iii (log2 fold change ≥1).

## 4. Discussion

In this study, we utilized our previously developed LC-MS based non-targeted metabolomics method to analyze the extracellular metabolites of hiPSC_585A1 cells under three different culturing conditions: 1) hiPSC medium consisting of mTeSR plus, 2) the E6 basal differentiation medium, and 3) the E6 basal medium combined with chemical inhibitors and growth factors to promote differentiation towards ectodermal or mesodermal lineages. The xeno-/serum-free E6 basal medium is designed to support differentiation with a formulation of nutrients – that will be discussed later–while lacking essential components such as TGF-β and bFGF, which are required for hiPSC survival and self-renewal. This allows for flexible control over differentiation without compromising other essential components^26^. We used two types of inhibitors for Wnt/ β-catenin inhibition, IWP-2 and IWR-1 endo which have been extensively reported in many differentiation protocols, involving neuron cells ^27, 28^, cardiomyocytes ^5^, and retinal and corneal epithelial cells cells and two types for the inhibition of TGF-β kinase/activin receptor that are commonly reported in the differentiation of the forehead and eye lineage SB505124 ^29^, and A83-01^30^, where the inhibition potency of A83-01 is much greater than that of other TGF-β/R inhibitors ^31^. Previously we reported the combination of 2.5 µM of A83-01 with IWR-1 endo and the growth factor bFGF for the initiation of the differentiation toward the corneal epithelium lineage which is initially diverted from the surface ectoderm^6^. Moreover, we used CHIR99021 as a potent inhibitor of glycogen kinase 3 (GSK-3) that is commonly used with hPSCs for the diversion into mesodermal lineage prior to the cell differentiation into cardiomyocytes or myoblasts ^32, 33^. Additionally, we used Y-27632 as Rho associated coiled coil kinase (ROCK) inhibitor that is frequently used with hPSCs for the enhancement of their survival ^34^.

To confirm the differentiation process in the samples under E6 only or in combination with different treatment, we conducted immunofluorescence staining of the pluripotency marker OCT3/4. Our results showed a significant reduction of the OCT3/4 expression in all samples that were cultured under E6 only or in combination with chemicals or the growth factor bFGF as compared with cells under the mTeSR plus medium which indicated the initiation of the differentiation in cells. In fact, the use of E6 only has been reported to divert the cells toward the neural crest progenitors that are derived from the ectodermal lineage^26^ and the combination of E6 with bFGF was also used to generated myoblast cells that are diverted from mesodermal lineage^35^. Further combination with A83-01, IWP-2 _ SB505124_ bFGF, IWP-2_A83-01_ bFGF, and IWR-1 endo_ SB505124 led to greater reduction in the OCT3/4 indicating more potency in the initiation of the differentiation of cells. We have previously reported that the inhibition of Wnt/ β-catenin and TGF-β kinase/activin receptor in combination with bFGF can divert the cells toward the cornel lineage passing by the ectodermal lineage in the first week of differentiation where the eye ectodermal marker PAX6 is highly upregulated ^6^.

It is worth noting that we observed a separation in the PCA analysis between mTeSR plus and E6 medium at 0 hr, which indicated a difference in their composition. For example, our analysis revealed that mTeSR plus had a higher amount of alanyl-glutamine and alanyl-glycine compared to E6, which might explain the significant increase in glutamine and alanine in samples under mTeSR plus. This increase could be attributed to the gradual degradation of alanyl-glutamine over time. To accurately evaluate the changes in metabolites originating from cells under E6 and mTeSR plus conditions, we focused on metabolites that had similar levels at 0 hr or those that showed significant changes in abundance between 0 hr and 48 hrs. It has been reported that kynurenine secretion, which is a byproduct of tryptophan, can serve as a marker for pluripotency in embryonic stem cells ^36^. Decreased secretion of kynurenine has been linked to the initiation of differentiation towards the three germ layers, specifically the ectoderm. Our findings showed a significant decrease in kynurenine secretion between cells undergoing ectodermal differentiation as early as 48 hrs, which correlated with a decrease in tryptophan consumption.

The secretion of lactic acid is a hallmark of the glycolytic metabolism in cells, known as the Warburg effect, which is characteristic of both hPSCs and cancer cells and involves their dependence on glucose for energy ^37, 38^. However, we did not observe a significant difference in lactic acid levels between the groups under mTeSR plus medium or E6 medium after 48 hrs. This supports the findings of Yamamoto et *al*., who found a reduction in lactic acid secretion only after 4-5 days of differentiation^36^. This suggests that 48 hrs might not be a sufficient time frame to observe changes in lactic acid secretion.

It is well known that during hPSCs differentiation, cells transition from glycolysis to oxidative phosphorylation (OXPHOS) and the citric acid cycle, with pyruvic acid playing a crucial role in this process^39^. The addition of pyruvic acid to hPSCs has been reported to enhance differentiation towards the ectodermal and mesodermal lineages^40^. In this study, we observed a significant decrease in the level of exogenous pyruvic acid in cells under ectodermal differentiation conditions compared to the hiPSC cells under mTeSR plus medium conditions. This suggests that the initiated OXPHOS metabolism in differentiated cells may have consumed the pyruvic acid. On the other hand, the low consumption of pyruvic acid in hiPSCs is likely due to the active glycolysis process that generates endogenous pyruvic acid, which compensates for the consumption of exogenous pyruvic acid.

This is the first study utilizing metabolomic profiling of micro-scaled cell culture medium (CCM) from human induced pluripotent stem cells (hiPSCs) during their early differentiation. We were able to detect the initiation of differentiation by observing changes in key biological metabolites such as glucose, pyruvic acid, and lactic acid, as well as in the markers of hiPSCs metabolites, kynurenine, and tryptophan. Additionally, a group of metabolites that tend to be specific to ectodermal differentiation were noticed, such as the secretion of indole-3-lactic acid, oxalic acid, and adenosine, and the decrease of 2-oxoglutaric acid. Further research is needed to determine the significance of these discovered metabolites in the differentiation towards specific lineages.

## Acknowledgments

We acknowledge the Ritsumeikan Global Innovation Research Organization (R-GIRO) and the Gunma University Initiative for Advanced Research (GIAR) for their support. The funding for this work was generously provided by the Japan Society for the Promotion of Science (JSPS) through grants 20K20168, 22K14548 and 20KK0160, as well as by the Hirose Foundation to Rodi Abdalkader.

## Authors Contributions

R.A. was responsible for the overall research concept, project management, design and execution of biological experiments, data analysis and interpretation, visualization of results and writing of the manuscript. R.C. performed the non-targeted metabolomic LC-MS analysis, contributed to data interpretation and editing of the manuscript. T.F. provided insight on data interpretation. All authors critically reviewed the manuscript prior to publication and gave their approval.

## Supplementary information

**Figure S1.**
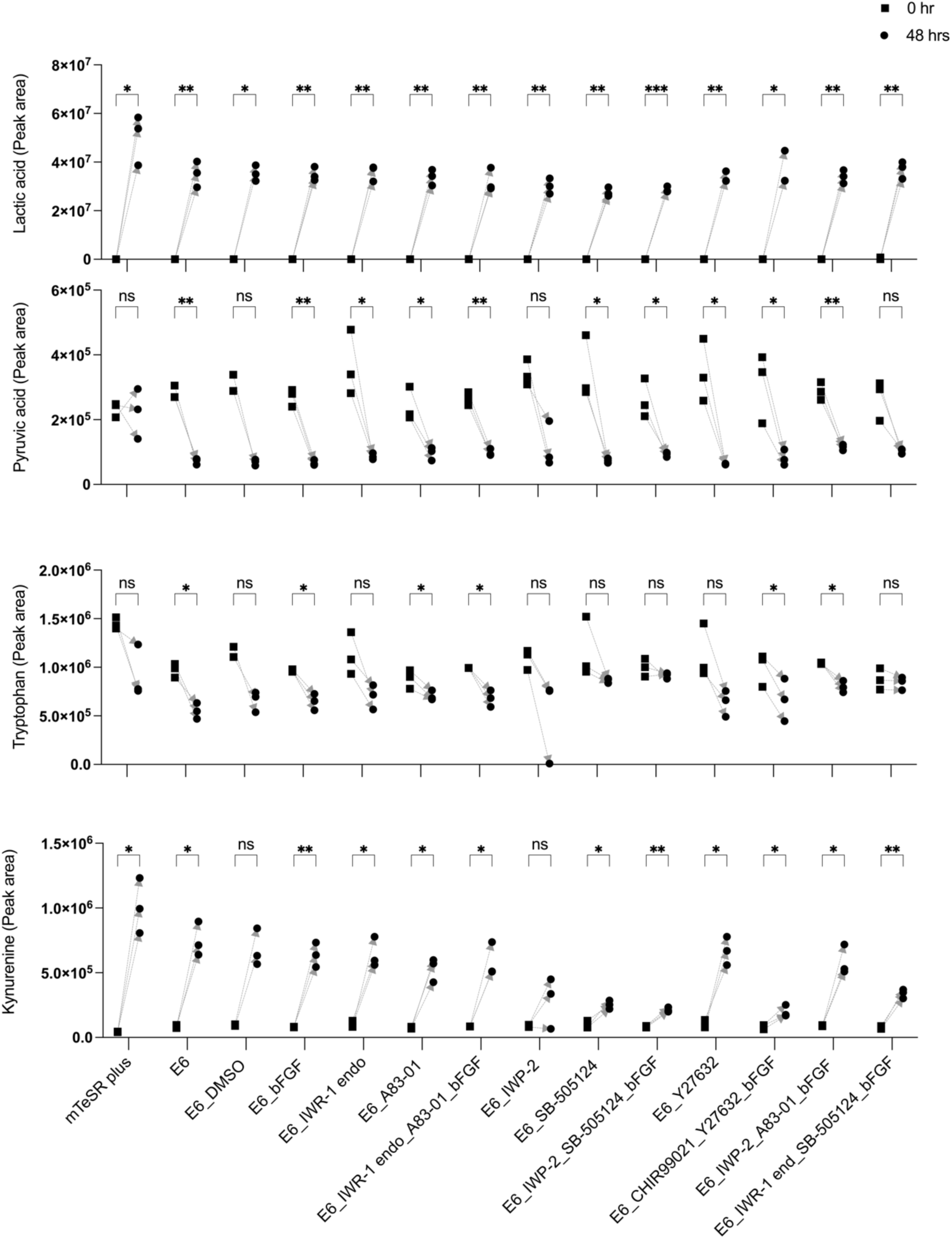
Analysis of extracellular metabolites from the human iPSC-585A1 at 0 hr and 48 hrs. Representative examples of selected metabolites including: lactic acid, pyruvic acid, tryptophan, and kynurenine. The p-values were determined using a paired t-test, with a significance level of * p <0.05, ** p <0.01 and *** p <0.001. No significant difference was noted for metabolites with a p-value of “ns”.

**Figure S2.**
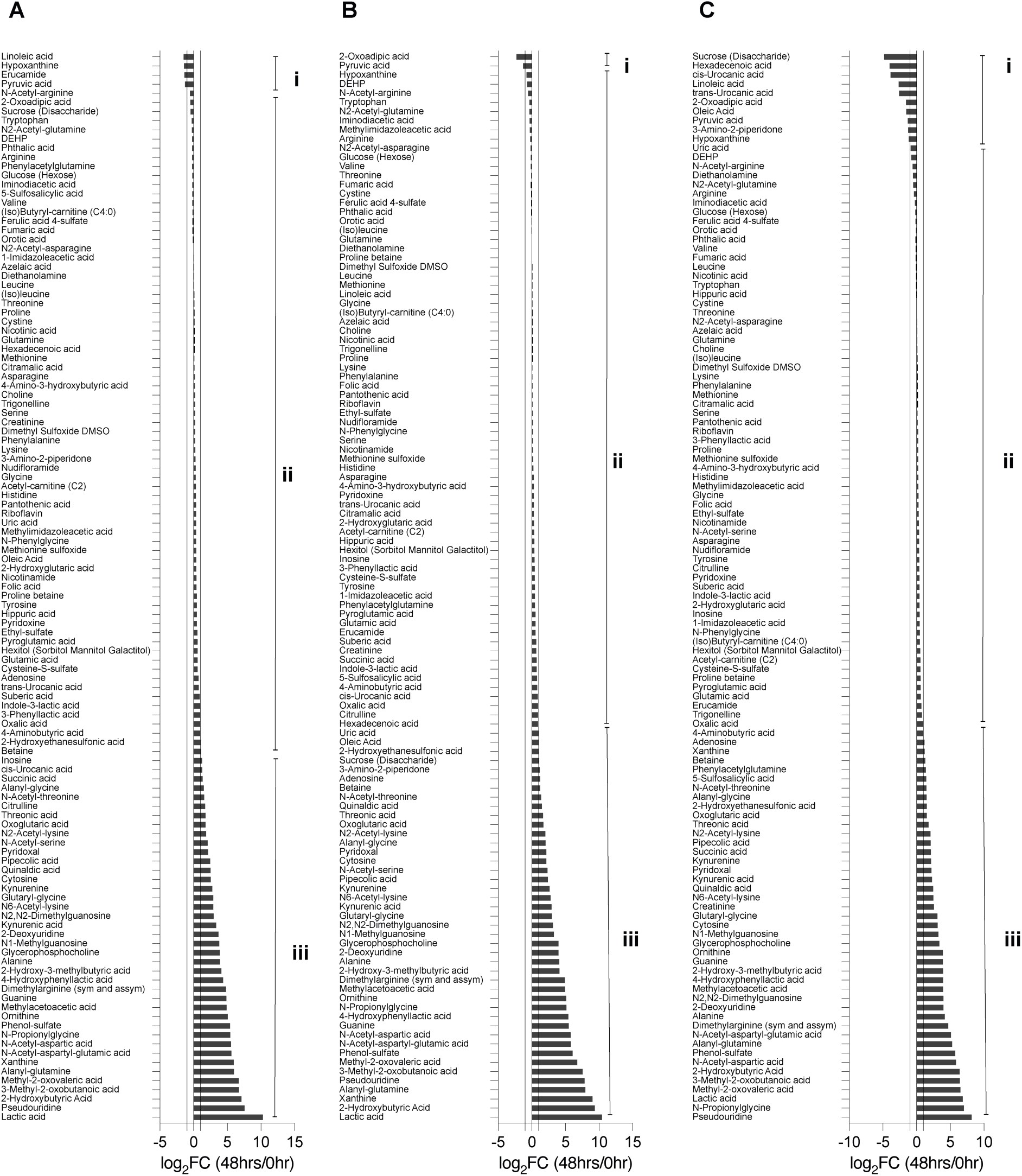
The determination of metabolites changes after 48 hrs. The ratio folds change (log_2_ 48 hrs/0 hr) of metabolites abundances. (A) E6 in combination with A83-01(B). E6 in combination with IWP-2 _ A83-01_ bFGF. (C) E6 in combination with IWR-1 endo _ SB505124_ bFGF. Category i (log2 fold change ≤-1), Category ii (log2>-1 and log2 < 1), and Category iii (log2 fold change ≥1.

**Figure S3.**
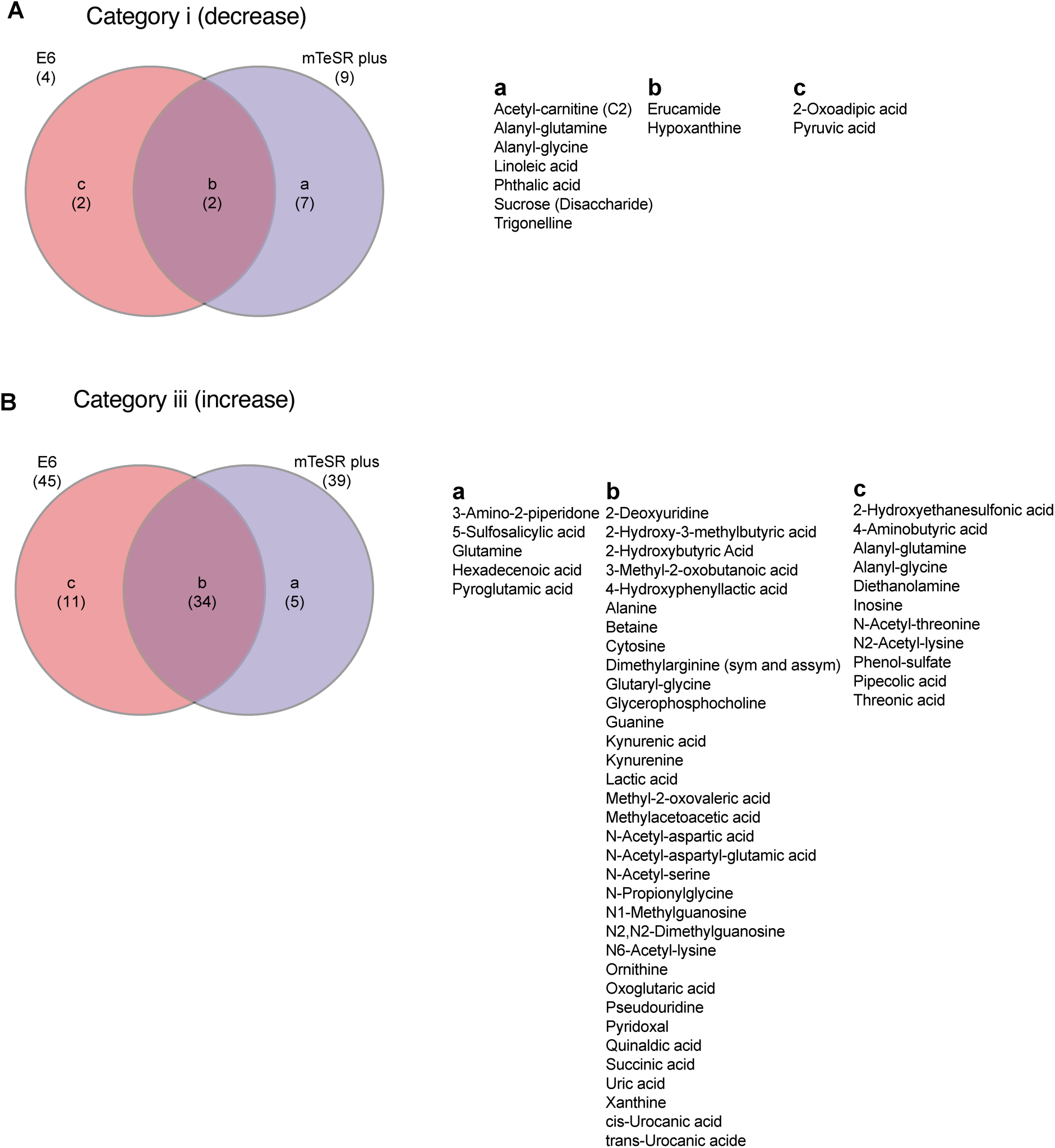
The determination of metabolites signatures under the differentiation conditions. Venn diagrams that highlight the differential shifts of metabolites between the hiPSC medium (mTeSR plus) and the differentiation medium E6. (A) In Category i (log2 fold change ≤-1). In Category iii (log2 fold change ≥1).

## References

1. Thomson, J. A. Embryonic stem cell lines derived from human blastocysts. Science (*80*-.). 282, 1145–1147 (1998).

2. Takahashi, K. & Yamanaka, S. Induction of pluripotent stem cells from mouse embryonic and adult Fibroblast cultures by defined factors. Cell 126, 663–676 (2006).

3. Greenhough, S., Medine, C. N. & Hay, D. C. Pluripotent stem cell derived hepatocyte like cells and their potential in toxicity screening. Toxicology vol. 278 250–255 (2010).

4. Zhang, R. et al. Identification of proliferating human hepatic cells from human induced pluripotent stem cells. Transplant. Proc. 46, 1201–1204 (2014).

5. Lian, X. et al. Directed cardiomyocyte differentiation from human pluripotent stem cells by modulating Wnt/β-catenin signaling under fully defined conditions. Nat. Protoc. 8, 162–175 (2013).

6. Abdalkader, R. & Kamei, K. An efficient simplified method for the generation of corneal epithelial cells from human pluripotent stem cells. Hum. Cell. 35, 1016–1029 (2022).

7. Susaimanickam, P. J. et al. Generating minicorneal organoids from human induced pluripotent stem cells. Dev. 144, 2338–2351 (2017).

8. Shimada, H., Yoshimura, N., Tsuji, A. & Kunisada, T. Differentiation of dopaminergic neurons from human embryonic stem cells:modulation of differentiation by FGF-20. J. Biosci. Bioeng. 107, 447–454 (2009).

9. Rajasingh, S. et al. Comparative analysis of human induced pluripotent stem cell-derived mesenchymal stem cells and umbilical cord mesenchymal stem cells. J. Cell. Mol. Med. 25, 8904–8919 (2021).

10. Guijas, C., Montenegro-Burke, J. R., Warth, B., Spilker, M. E. & Siuzdak, G. Metabolomics activity screening for identifying metabolites that modulate phenotype. Nature Biotechnology. 36, 316–320 (2018).

11. Dong S., Yan, Z. & Yang, H. A Sensitive precolumn derivatization method for determination of piperazine in vortioxetine hydrobromide using a C8 column and high-performance liquid chromatography-mass spectrometry. Anal. Sci. 32, 1333–1338 (2016).

12. Naz, S., Moreira dos Santos, D. C., García, A. & Barbas, C. Analytical protocols based on LC–MS, GC–MS and CE–MS for nontargeted metabolomics of biological tissues. Bioanalysis 6, 1657–1677 (2014).

13. Takashina, S., Igarashi, Y., Takahashi, M., Kondo, Y. & Inoue, K. Screening method for the quality evaluation of cannabidiols in water-based products using liquid chromatography tandem mass spectrometry. Anal. Sci. 36, 1427–1430 (2020).

14. Abdalkader, R. et al. Untargeted LC-MS metabolomics for the analysis of micro-scaled extracellular metabolites from hepatocytes. Anal. Sci. 37, 1049–1052 (2021).

15. Abdalkader, R., Chaleckis, R., Wheelock, C. E. & Kamei, K. Spatiotemporal determination of metabolite activities in the corneal epithelium on a chip. Exp. Eye Res. 209, 108646 (2021).

16. Okita, K. et al. An efficient nonviral method to generate integration-free human-induced pluripotent stem cells from cord blood and peripheral blood cells. Stem Cells 31, 458–466 (2013).

17. Naz, S. et al. Development of a liquid chromatography–high resolution mass spectrometry metabolomics method with high specificity for metabolite identification using all ion fragmentation acquisition. Anal. Chem. 89, 7933–7942 (2017).

18. Chaleckis, R., Naz, S., Meister, I. & Wheelock, C. E. LC-MS-based metabolomics of biofluids using all-ion fragmentation (AIF) acquisition. Methods in molecular biology. 1730, 45–58 (2018).

19. Tada, I. et al. Creating a reliable mass spectral–retention time library for all ion fragmentation-based metabolomics. Metabolites 9, 251 (2019).

20. Chambers, M. C. et al. A cross-platform toolkit for mass spectrometry and proteomics. Nat. Biotechnol. 30, 918–920 (2012).

21. Tsugawa, H. et al. MS-DIAL: Data-independent MS/MS deconvolution for comprehensive metabolome analysis. Nat. Methods 12, 523–526 (2015).

22. Tsugawa, H. et al. A lipidome atlas in MS-DIAL 4. Nat. Biotechnol. 38, (2020).

23. McQuin, C. et al. CellProfiler 3.0: Next-generation image processing for biology. PLOS Biol. 16, e2005970 (2018).

24. Chong, J. et al. MetaboAnalyst 4.0: Towards more transparent and integrative metabolomics analysis. Nucleic Acids Res. 46, W486–W494 (2018).

25. Sumner, L. W. et al. Proposed minimum reporting standards for chemical analysis: chemical analysis working group (CAWG) metabolomics standards initiative (MSI). Metabolomics 3, 211–221 (2007).

26. Lippmann, E. S., Estevez-Silva, M. C. & Ashton, R. S. Defined human pluripotent stem cell culture enables highly efficient neuroepithelium derivation without small molecule inhibitors. Stem Cells 32, 1032–1042 (2014).

27. Oosterveen, T. et al. Pluripotent stem cell derived dopaminergic subpopulations model the selective neuron degeneration in Parkinson’s disease. Stem Cell Reports 16, 2718– 2735 (2021).

28. Muckom, R. et al. High-throughput 3D screening for differentiation of hPSC-derived cell therapy candidates. Sci. Adv. 6, eaaz1457 (2020).

29. Takamiya, M. et al. Pax6 organizes the anterior eye segment by guiding two distinct neural crest waves. PLOS Genet. 16, e1008774 (2020).

30. Theerakittayakorn, K. et al. Differentiation induction of human stem cells for corneal epithelial regeneration. Int. J. Mol. Sci. 21, 7834 (2020).

31. Tojo, M. et al. The ALK-5 inhibitor A-83-01 inhibits Smad signaling and epithelial-to-mesenchymal transition by transforming growth factor-β. Cancer Sci. 96, 791–800 (2005).

32. Kempf, H. et al. Controlling expansion and cardiomyogenic differentiation of human pluripotent stem cells in scalable suspension culture. Stem Cell Reports 3, 1132–1146 (2014).

33. Shelton, M. et al. Derivation and expansion of PAX7-positive muscle progenitors from human and mouse embryonic stem cells. Stem Cell Reports 3, 516–529 (2014).

34. Gauthaman, K., Fong, C. Y. & Bongso, A. Effect of ROCK inhibitor Y-27632 on normal and variant human embryonic stem cells (hESCs) in vitro: its benefits in hESC expansion. Stem Cell Rev. Reports 6, 86–95 (2010).

35. Xuan, W., Khan, M. & Ashraf, M. Pluripotent stem cell-induced skeletal muscle progenitor cells with givinostat promote myoangiogenesis and restore dystrophin in injured Duchenne dystrophic muscle. Stem Cell Res. Ther. 12, 131 (2021).

36. Yamamoto, T. et al. Kynurenine signaling through the aryl hydrocarbon receptor maintains the undifferentiated state of human embryonic stem cells. Sci. Signal. 12, eaaw3306 (2019).

37. Warburg, O., Wind, F. & Negelein, E. The metabolism of tumors in the body. J. Gen. Physiol. 8, 519–30 (1927).

38. Ito, K. & Suda, T. Metabolic requirements for the maintenance of self-renewing stem cells. Nat. Rev. Mol. Cell Biol. 15, 243–256 (2014).

39. Zhang, J., Nuebel, E., Daley, G. Q., Koehler, C. M. & Teitell, M. A. Metabolic regulation in pluripotent stem cells during reprogramming and self-renewal. Cell Stem Cell 11, 589–595 (2012).

40. Song, C. et al. Elevated exogenous pyruvate potentiates mesodermal differentiation through metabolic modulation and AMPK/mTOR pathway in human embryonic stem cells. Stem Cell Reports 13, 338–351 (2019).

